# Effects of drying and milling treatments on estimates of foliar functional traits with VSWIR spectroscopy

**DOI:** 10.64898/2026.11.11.765977

**Authors:** Zachary Breuer, Annabelle Majerus, Henry A. Frye, Philip A. Townsend

## Abstract

Spectroscopy of dried, ground foliage is a cost-efficient measurement to estimate leaf biochemical traits in agricultural and ecological research. Drying removes water that obscures biochemical absorption features, and milling homogenizes samples, yet how drying and milling methods affect spectra and trait predictions is understudied. We compared two drying methods (oven-drying; flash-freezing then freeze-drying) and three milling methods (coarse and fine rotary milling; ball milling) using 61 individuals of 11 broadleaf and needle-or scale-leaved woody species. Mean reflectance differences between milling treatments ranged 1.7–4.6% across the full spectrum and were largest between fine and coarse rotary milling. Freeze-dried samples had on average 7.2% higher reflectance than oven-dried samples, with larger differences in broadleaf species. Bayesian ANOVAs of spectrally predicted nitrogen and calcium demonstrated that drying had the largest effect, and that fine and ball milling also shifted predictions relative to coarse milling, but all treatment effects were smaller than interspecific variation. Although the partial least squares regression models were trained on oven-dried material, freeze-dried and more finely milled samples had lower prediction error (NRMSE) than oven-dried samples. Our finding emphasize that sample processing methods should be carefully documented for studies measuring traits from foliar spectroscopy.

**HIGHLIGHT STATEMENT:** Drying and milling methods impact the reflectance and subsequent trait predictions of foliar powders. Researchers should aggregate datasets with similar handling procedures for foliar trait remote sensing.

## INTRODUCTION

The biochemistry and morphology of plant foliage (collectively: foliar traits) impact individual fitness, and when upscaled across individuals in a community, drive ecosystem processes. A non-exhaustive list of such biochemical traits includes the concentrations of photosynthetic pigments, non-structural carbohydrates (sugars and starches), defense compounds, cell wall components, and the overall elemental and isotopic composition of leaves that are related to molecules that regulate physiological function (e.g., enzymes) or are the outcomes of photosynthesis and subsequent chemical pathways. Because of their ease of measurement, plant trait ecology has primarily focused on standardized structural traits such as leaf mass per area (Green *et al*., 2022) and only a small subset of chemical traits, e.g., foliar nitrogen and phosphorus, that fall along well-established trait-trait tradeoff axes (Wright *et al*., 2004; Díaz *et al*., 2016). However, a more expansive set of leaf chemical traits offers the potential of refining our understanding of the adaptive significance of traits, the role of evolutionary history in shaping patterns of traits, and how these traits link to ecosystem function (Thompson *et al*., 1997; Watanabe *et al*., 2007; Blumstein *et al*., 2023; Asao *et al*., 2025). A barrier to measuring these foliar chemical traits is that chemical assays are costly, labor-intensive, and require a high quantity of collected material when multiple chemical assays are conducted concurrently.

One approach that reduces these barriers is the use of spectral reflectance on dried and homogenized leaf material to predict multiple leaf biochemical traits (Norris *et al*., 1976; Serbin *et al*., 2014). Dried-leaf spectroscopy measures absorption features derived from the stretching of chemical bonds in the infrared region of the spectrum (1000–2500 nm) caused by a broad suite of plant compounds (Curran, 1989). By collecting an initial calibration dataset of foliar reflectance spectra and associated chemical traits, data-driven models such as partial least squares regression can be developed to estimate multiple chemistry values from a single spectral measurement (Foley *et al*., 1998; Wold *et al*., 2001; Burnett *et al*., 2021). This approach produces estimates of foliar chemistry at a fraction of the time and cost of direct laboratory measurement. It also can enable standardized approaches to trait estimations, given a common set of laboratory procedures. Early development of this technique was aimed at estimating the forage quality of plants for agricultural purposes (Norris *et al*., 1976) and since then, dried foliar spectroscopy has been shown to robustly estimate a large suite of plant chemical traits specifically for both ecological and agricultural applications (Bian *et al*., 2013; Serbin *et al*., 2014; Couture *et al*., 2016; Prananto *et al*., 2020; Wang *et al*., 2020). These measurements and their subsequent foliar trait predictions are now being used to generate validation chemistry data for trait mapping efforts from airborne and satellite-based image spectroscopy (Singh *et al*., 2015; Wang *et al*., 2020; Chlus and Townsend, 2022; Liu *et al*., 2023, 2024).

As satellite-based image spectroscopy progresses toward the goal of producing global foliar trait maps, there is a need to calibrate and validate predicted trait maps using data representing the range of ecosystems globally. Lab-measured data will be central to this effort, but it will be impractical to comprehensively cover Earth’s ecosystems - including phenological and ontogenetic variation – across a wide number of traits. Globally generalizable foliar spectroscopy models will bridge this gap by providing the necessary volume of trait estimates. This will require the ability to merge foliar trait-spectral datasets across data collection efforts, e.g., EcoSIS (Wagner *et al*., 2019). However, one impediment to this goal is the impact that different lab handling and sample processing procedures may have on foliar spectral measurements. If sample handling has a large impact on spectral measurements, this would be problematic not only for satellite validation efforts, but also for attempts at making ecological inferences from such data since different ecological scales, e.g., intra-versus inter-specific variation, could be conflated with sample handling differences. Despite the reliance on dried foliar spectroscopy in ecological remote sensing, little explicit consideration has been given to the differing methods of sample handling. Our goal is to address variations resulting from differing sample handling approaches; note that we do not address differences in laboratory chemical analyses, e.g., Quentin *et al*., (2015), which is a separate and significant issue that also has important implications.

To prevent decomposition and preserve tissue chemistry, leaf tissue samples are often oven-dried at low temperatures between 65–75°C (Bolster *et al*., 1996) or flash-frozen and then freeze-dried (Smith, 1973; Popp *et al*., 1996). While flash-freezing material may be the best method to preserve tissue chemistry, it is often impractical to access or transport liquid nitrogen or frozen CO_2_ (dry ice) to remote field locations; therefore, short-term air drying followed by oven-drying is often favored. Most previous remote sensing studies have used spectral measurements made on material that was oven-dried (Peterson *et al*., 1988; Bian *et al*., 2013; Singh *et al*., 2015; Lepine *et al*., 2016; Wang *et al*., 2020), though some have used frozen and freeze-dried material (Curran *et al*., 2001) or a hybrid approach where the freeze-dried material was subsequently oven-dried (Chlus and Townsend, 2022). Previous research in forage quality has assessed how drying treatments impact spectral measurements and trait predictions (Alomar *et al*., 1999a,b; Park *et al*., 2002; Ikoyi and Younge, 2020), but no comparison to date has been made on foliar material collected for the purposes of ecological study across a wider set of species. In addition to the drying method, how the foliar samples are homogenized via milling has also only been explored in the forage quality literature (Alomar *et al*., 1999b; Ikoyi and Younge, 2020) and remains unexamined in the ecological remote sensing context. The mesh size (screen opening) used to sieve and homogenize material may vary significantly causing visible differences in the foliar powder (Fig. 1). The earliest study often credited with founding the field of trait estimation from dried foliage used a mix of different grind sizes (Norris *et al*., 1976) and noted that differences in particle size do affect measurements.

**Figure 1:**
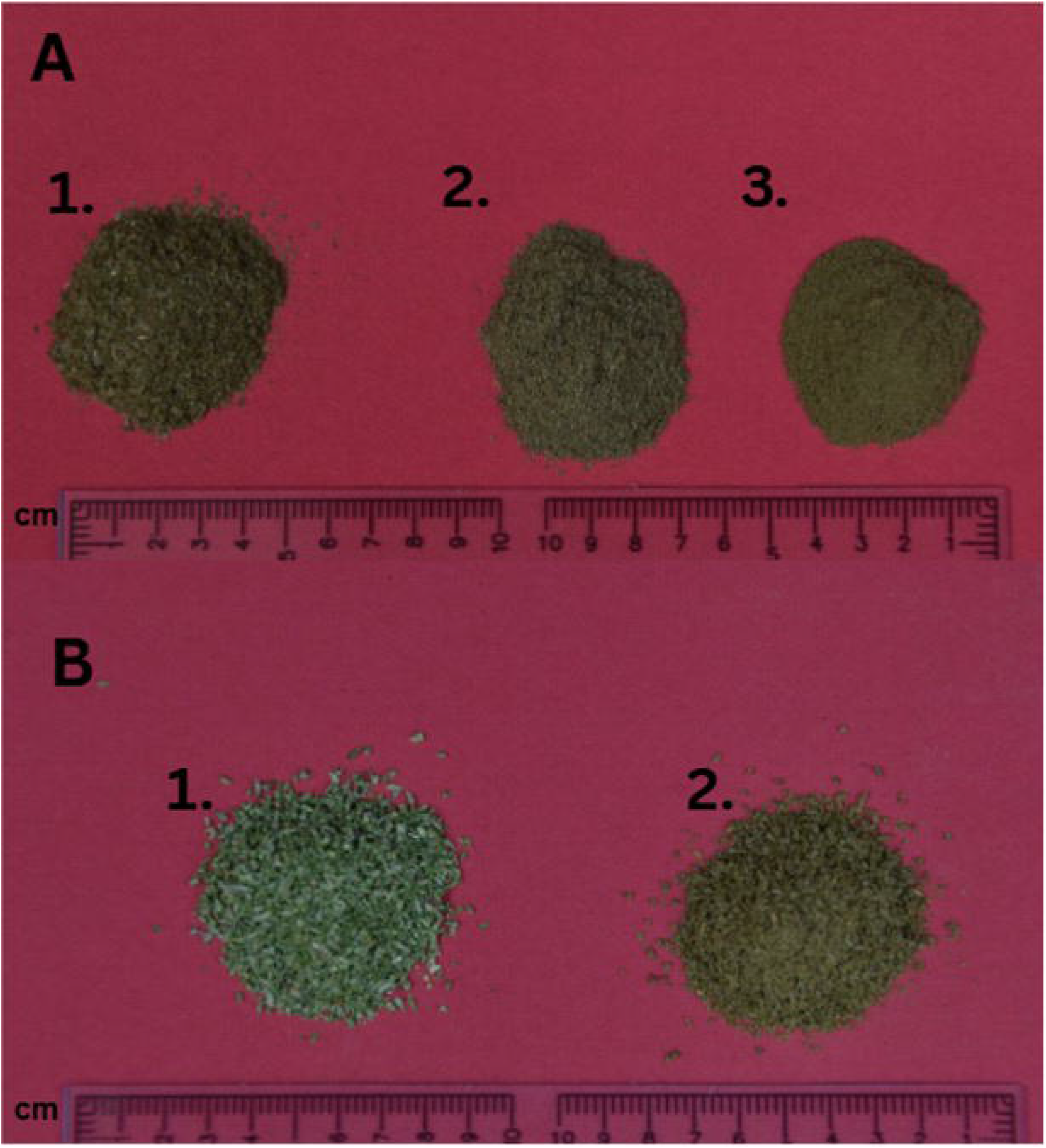
Milling and drying visibly change the color and texture of foliar powders. (A) Oven-dried Acer saccharum milled through *(1) a 20-mesh (850 µm) screen, (2) a ball mill and (3) a 60-mesh (250 µm) screen. (B)* Juniperus virginiana *that was (1) freeze-dried or (2) oven-dried. Rulers are in cm*. ***Alt Text Figure 1:*** *Photograph of foliar powders piled on a red background above centimeter rulers. In panel A, three piles of sugar maple powder run left to right from coarse and flaky (20-mesh), to intermediate (ball-milled), to fine and uniform (60-mesh), with texture becoming smoother as particle size decreases. In panel B, two piles of eastern red cedar powder: the freeze-dried pile on the left is pale green and the oven-dried pile on the right is darker olive brown*.

The current study examines the impact that foliar sample drying and milling have on spectral measurements and subsequent foliar trait predictions. Milling of foliar material is often conducted using two methods: either a rotary mill that grinds material by forcing it through a mesh screen (Wang *et al*., 2020) or a ball-mill where the impact of ball bearing media crushes the material into a powder (Kothari *et al*., 2024). In the case of the rotary mill, the particle size of the foliar powder is determined by the size of the mesh screen, with 20 (850 μm) or 60 (250 μm) mesh being the most used. Particle size in ball milling is in part determined by the duration that the material is left in the mill, with longer durations resulting in finer powders. However, increased duration in a ball mill can create frictional heat in the media and lead to degradation of temperature-sensitive leaf compounds; as well, differences in leaf morphology and composition (e.g., thick leaves, presence of hairs or latex) can also influence final particle size. The physical properties of the powder, e.g., surface area and gaps, vary between different milling procedures and have been shown to have significant impacts on spectral measurements (Norris *et al*., 1976; Myers *et al*., 2015).

This study explores the variation of spectral reflectance between these processing factors and assesses the impacts on the prediction of two foliar elements: nitrogen and calcium. Foliar nitrogen has been measured in ecological remote sensing applications for decades, and has strong model prediction generalizability across different image collection efforts (Wessman *et al*., 1988; Matson *et al*., 1994; Townsend *et al*., 2003; Martin *et al*., 2008; Kokaly *et al*., 2009). Statistical models typically leverage a number of wavelength features related to protein content across the shortwave infrared spectrum for nitrogen prediction (Curran, 1989). In contrast, spectroscopy predictions from calcium are more modest (Chadwick and Asner, 2016; Chlus and Townsend, 2022; Kothari *et al*., 2023) and are likely based on narrow features caused by calcium storage compounds in vacuoles such as oxalates (Cloutis, 2015), or co-correlated with traits related to cell wall composition and growth rates (Hawkesford *et al*., 2023).

To ensure the accuracy and effectiveness of foliar spectroscopy across different handling procedures, our study uses a factorial experiment to investigate different aspects of the sample preparation process before spectroscopy measurements. Sixty-one samples from both broad-leaf deciduous and evergreen needle- and scale-leaf trees were collected and then either oven-dried or flash-frozen. These samples were then further split into three milling treatments: 1) a ball-mill 2) a rotary mill at a coarse mesh size or 3) a rotary mill at a fine mesh size. We examine the effect that each treatment has on predictions of nitrogen and calcium using previously published partial least squares regression coefficients (Wang *et al*., 2020) and evaluate whether sample processing variation is greater than the observed intra- and inter-specific variation.

## MATERIALS AND METHODS

### Sample Collection

Foliar samples were collected from 11 species in the Lakeshore Nature Preserve at the University of Wisconsin-Madison from late July to August 2024 (Supplementary Fig. S1). Tree species were sampled if their canopy was sunlit and not overtopped by other trees. Pole pruners and an Air Powered Tree Access launcher (TreeStuff, Greensboro, NC, USA) modified with a fishing reel were used to reach sunlit branches, with the latter used for tall canopies. Samples from four deciduous broadleaf species and two deciduous understory shrubs (5 individual trees per species, for a total of 30 individuals) were collected from sugar maple (*Acer saccharum),* red oak *(Quercus rubra),* black locust (*Robinia pseudoacacia),* hackberry (*Celtis occidentalis),* and the shrubs buckthorn (*Rhamnus cathartica)* and pagoda dogwood (*Cornus alternifolia)*. With one exception, five species from evergreen needle or scale-leaved trees (6 individual trees per species) were collected from red pine (*Pinus resinosa),* Norway spruce (*Picea abies)*, red cedar (*Juniperus virginiana)*, white pine (*Pinus strobus*; had a total 7 individual collections), and white cedar (*Thuja occidentalis).* A total of 31 individuals was collected for the evergreen needle or scale-leaved trees. Leaves from whole branches from each tree were kept intact and processed within 3–4 hours of collection.

### Sample Processing

Once collected, broad leaves were separated from the petiole so that only the leaf lamina was included in spectral and trait measurements. Needle leaves were separated above the fascicle bundles to avoid non-photosynthetic tissue. Processed leaves were split into six treatments: flash frozen/freeze-dried and ball-milled, flash frozen/freeze-dried and milled with a rotary mill with a fine mesh, flash frozen/freeze-dried and milled with a rotary mill with a coarse mesh, oven-dried and ball-milled, oven-dried and milled with a rotary mill with a fine mesh, oven-dried and milled with a rotary mill with a coarse mesh. Each treatment per sample was weighed to a minimum of 5 g to ensure adequate material quantity for spectral and chemical measurements.

Oven-dried samples were placed in a forced-air oven (IsoTemp Oven, Fisher Scientific, Waltham, MA, USA) for 48–72 hours at 65° C until samples were dried. Samples for the freeze-dried treatment were flash-frozen as soon as leaf material was separated from the main branch by submerging paper coin envelopes containing the leaf material in liquid nitrogen. After flash-freezing, samples were then placed in a freeze dryer (24DX48 Specimen Freeze Dryer, Virtis Company, Gardiner, NY, USA) until samples were dried without any visible signs of moisture, e.g., dark spots.

Once dried, all samples were milled through an E3300 Eberbach Corporation (Belleville, MI, USA) single-speed rotary mill using a 20-mesh (850 μm; the coarse mesh;. A subset of the samples was further milled with a ball mill or using a fine mesh (40-or 60-mesh; 425 and 250 μm, respectively) with the rotary mill. For the fine mesh rotary treatment, all broadleaf samples were ground on a step-wise basis, where each sample was first ground to 20-mesh and then further ground to the specification of labeling (e.g. 60-mesh samples were ground to 20-mesh, then 40-mesh, and finally 60-mesh). Unlike the broadleaf samples, needle-leaf samples were milled to 40 mesh instead of 60-mesh due to difficulty in milling samples with high resin content to a finer particle size. The ball milling treatment followed protocols documented by Frye *et al*., (2026c) where coarsely ground samples were placed in 20 ml plastic scintillation vials along with thirty 3 mm steel ball bearings. Our approach deviated from the published protocol in that one 9 mm bearing was also added to prevent samples from clumping with the smaller ball media. Vials were then placed into a modified paint shaker (C1-10038 Paint Shaker, Fleming Gray Limited, St. George, Ontario, Canada) for 1 minute, rotated 180 degrees, and then shaken for another minute (for a total of 2 minutes). Ball bearings were extracted from the vials with a magnet and then cleaned with water, followed by an acetone rinse to prevent rust.

### Spectral measurements, trait predictions, and chemistry validation

Spectroscopy measurements (350–2500 nm) of ground foliar material were made with an ASD FieldSpec 3 (Analytical Spectral Devices, Boulder, Colorado, USA) and ASD plant-probe fore optic mounted onto a custom-built press apparatus (Serbin *et al*., 2014; Frye *et al*., 2026a). Prior to measurement, each sample was packed into a 3-D printed cup consisting of three wells with an approximate depth of 6.5 mm. After being packed, oven-dried material was placed in a low-temperature oven (65 °C) overnight to remove any residual moisture that may have accrued during milling and packing. Samples were then kept in a glass jar desiccator with silica packets while awaiting spectral measurement. Freeze dried samples were placed in a freeze drier for a minimum of 48 hours following packing to eliminate any moisture gained during milling and packing. After being removed from the freeze drier, packed samples were kept at cold temperatures in sealed refrigeration units until spectral measurements were performed. While samples were queued for spectral measurements, packed samples were kept in coolers with cold packs to ensure minimal moisture gain. Sample temperature and humidity at the time of scanning were not recorded, and the freeze-dried samples were not deliberately equilibrated to room temperature before measurement. Each well is measured and rotated 180° to account for anisotropy in the material, resulting in a total of six spectral replicates per sample. Spectra were jump-corrected by an additive offset at the 1000 and 1800 nm detector splices holding the 1001–1800 nm detector fixed using code from Kovach (2023).

To assess the impact that different processing treatments would have on trait predictions, a subset of the oven-dried rotary-milled (20-mesh) material was sent to Brookside Laboratories (New Bremen, OH, USA) for elemental analysis. Nitrogen was measured via combustion with the Dumas method, and calcium was measured with inductively coupled plasma optical emission spectroscopy (ICP-OES). Other elemental concentrations were measured with ICP-OES and are provided by Majerus *et al*., (2026), but are not the focus of the current study. Given that only one treatment of material was assayed directly, we cannot assume that this reflects each treatment’s accuracy. It is possible that the N concentration is overestimated for the oven-dried material, as drying may lead to carbon volatilization, thus increasing the relative mass of N in a sample (Mayland, 1968). Some carbon volatilization may also occur due to vacuum pressure in a freeze-drier, though the volatilization is likely still less than oven-drying (Jones and O’Hara, 2016). As a *post hoc* check of these assumptions, we measured 11 paired oven-dried and freeze-dried (both ball-milled) samples for nitrogen and carbon with isotope-ratio mass spectrometry (IRMS). IRMS measures were used because this could be done at a local facility and perturbations to sample material due to the transport would be kept to a minimum. These values (along with isotopic compositions) were calculated based on the modified Dumas combustion method using a FlashSmart Elemental Analyzer (ThermoScientific, Waltham, MA, USA) coupled to a Delta Q IRMS (ThermoScientific, Waltham, MA, USA). Precision of elemental composition measurements was ±1.2% for total carbon (%C) and ±0.07% for total nitrogen (%N) based on percentage points of dry mass (one standard deviation of eight replicate analyses of NIST 1547).

### Particle size measurements

Following spectral measurements, a subset of 22 individuals (11 broadleaf, 11 needleleaf) was selected to examine the impacts of milling treatment on particle size. To investigate the difference in particle size between milling treatments, the size 20-mesh-milled, ball-milled, and finely-milled (40- and 60-mesh for needle and broad leaves respectively) samples were measured in an LS 13 320 Laser Diffraction Particle Size Analyzer (Beckman Coulter, Brea, CA, USA). The Particle Size Analyzer calculates a particle size distribution within a sample as described in Eshel *et al*., (2004). Distribution moments were averaged across growth form and milling treatment to summarize outputs.

### Statistical analysis

#### Analysis of variance

The variation across milling and sample drying treatments was partitioned as a function of spectrally predicted nitrogen and calcium values in two separate Bayesian ANOVA models. Milling treatment, sample drying treatment, and their interaction were treated as fixed effects. Random intercepts were included for species, individual tree within species, and tree by treatment combination; the six replicate scans of each tree under each milling × drying combination share the last term, and scan-to-scan variation is the residual. The brms formula was trait ∼ MillTreat * DryTreat + (1 | Species_Code) + (1 | Species_Code:SampleID) + (1 | Species_Code:SampleID:measure_id) with a Gaussian family. The fine mesh treatment included both broadleaves ground through 60-mesh sieve and needle-leaves through a 40-mesh sieve. Because growth form is nested within species, milling coefficients are averages across the two intensities and growth-form-specific milling effects were not estimated. Coefficient estimates were expressed in reference to samples that were oven-dried and milled with a rotary mill through a 20-mesh screen. Models were fitted in R 4.4.1 with the *brms* 2.23.0 package (Bürkner, 2017) and summarized using the *MCMCvis* package (Youngflesh, 2018), using four chains with 20,000 iterations and 5,000 warm-up. Coefficients had weakly informative Student-t priors (3 degrees of freedom, location 0, scale 2.5 mg g⁻¹). We kept brms defaults for the other parameters: intercept Student-t (3, 18.3, 5.7) for nitrogen and (3, 12.9, 9.4) for calcium, and half-Student-t (3, 0, 5.7) and (3, 0, 9.4) for the random-effect and residual SDs. R^ was ≤ 1.001 and bulk and tail effective sample sizes exceeded 10,900 for all fixed and group-level parameters, with no divergent transitions in 60,000 post-warm-up draws per model. In the calcium model, 37 transitions reached the maximum tree depth (12), which affects sampling efficiency but not validity. Percent changes are relative to the observed mean of the reference treatment (oven-dried, 20-mesh). Posterior predictive checks (overlays of the observed distribution and posterior-predictive draw; Supplementary Fig. S2) indicated that the models adequately reproduce the data. To partition variation among inter-specific, intra-specific, and treatment sources, we computed variance components from the posterior draws: the squared group-level standard deviations for species, tree within species, tree by treatment combination, and the residual (scan-to-scan) variance, and, for treatment, the variance across the six drying by milling cell means, including their interaction. Within each draw, each component was divided by the sum of all five (the total model variance) and summarized as the median and 95% interval; the ratio of the species standard deviation to the treatment standard deviation was computed per draw in the same way.

#### Spectral variation between treatments

To quantify treatment-level differences between spectral measurements, the difference spectrum was calculated as follows:

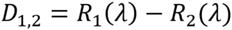

Where R_1_(λ) refers to the reflectance at a wavelength λ, measured under one milling treatment for a sample, and R_2_(λ) refers to the reflectance of a wavelength under another treatment for the same sample. Within the six replicates per sample, all possible pairwise difference spectra were calculated between two treatments for each sample. The mean of the distribution was used for sample statistics in figures and tables. To make inferences across treatments, the average spectral difference across all wavelengths was calculated for each pairwise comparison. For each tree, difference spectra were averaged over all 36 replicate pairs and over the wavelengths in each spectral region with the mean calculated across trees ± the half-width of the 95% confidence interval (t distribution, n = 61, 30 and 31 trees), treating trees as independent. All differences are absolute and expressed in percentage points of reflectance. To account for the fact that the finest milling treatment differs between broad- and needle-leaved species (the latter could only be ground through a 40-mesh screen), we built a linear regression across the 21 trees with particle size measures, regressing the particle size difference and a growth form term onto the overall differences in spectral differences between milling treatments The same analysis was repeated but with the spectral pre-processing step of vector-normalization (Feilhauer *et al*., 2010) which were how reflectance spectra were treated in the PLSR trait models from Wang *et al*., (2020). Vector-normalized (VN) spectra were calculated by dividing each reflectance by its full reflectance norm, i.e., 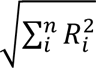 for band *i* to *n*.

Because the near infrared red spectroscopy literature often uses the standard normal variate (SNV) correction as a way to account for particle size differences in material (Barnes *et al*., 1989), we compared how spectral variability differed between SNV and VN spectra. For each normalization, the total sum of squares of the spectra between 1400 and 2450 nm (the range used for trait prediction, see next section) was partitioned into nested components: individual trees, dry treatment within trees, milling treatment within trees and drying treatment, and replicate scans. VN was calculated over either 350–2500 nm or 1400–2500 nm.

#### Trait prediction performance

Nitrogen and calcium were predicted using partial least squares regression (PLSR) coefficients reported in Wang *et al*., (2020) — a study that trained foliar models from species across the eastern US and thus covers the variation that we measured in our dataset. Samples in that study were oven-dried and milled using a rotary mill with a 20-mesh screen. To match the spectral pre-processing steps used by (Wang *et al*., 2020), spectra were unit vector-normalized (vn-spectra) and resampled to 5 nm resolution. Vector normalization serves a crucial role in the trait prediction process; vector normalization minimizes multiplicative differences in overall reflectance between spectra, e.g., illumination and particle-size related albedo, yielding more consistent trait predictions. Performance of predicted traits was assessed based on the root mean square error (RMSE) and normalized root mean squared error (NRMSE). RMSE and NRMSE were calculated as follows:

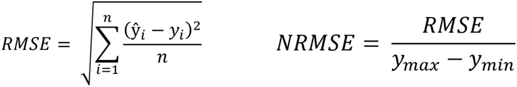

where, *ŷ* is the predicted value, *y* is the observed (Brookside) value, n = 61 trees per treatment, and *y_max_* − *y_min_* is the observed measured range.

In the chemometric literature, MSE (RMSE^2^) is often decomposed into two components: squared bias and residual variance (Bellon-Maurel *et al*., 2010). The bias is the mean offset between observed and predicted values (mean predicted – mean observed), whereas the residual variance conflates both deviation from a slope of 1 (proportionality) and precision (unsystematic error) around the relationship between the predicted and observed data. Because the residual variance conflates two different factors, we follow a three part decomposition of MSE: bias, proportionality, and precision (Willmott, 1981; Robeson and Willmott, 2023). We regressed predicted on observed values, calculating the proportionality component as (slope – 1)^2^ × the variance of observed values and the precision component is the mean squared residual about the regression line. To assess the overall agreement between predicted and reference values (oven-dried 20-mesh) for each treatment, the Bland-Altman bias and concordance correlation coefficient were calculated (Bland and Altman, 1986; Lin, 1989). The latter measure is decomposed into a precision factor (*ρ*) and accuracy term (Cb). Lastly, we calculated the offset of directly measured nitrogen and carbon values between freeze-dried and oven-dried material (freeze-dried – oven-dried). This gives an approximate bound on the degree to which model performance could be affected by the impact on chemical measurement from the drying treatment. To test whether the freeze-dried advantage depended on the oven-dried reference, we shifted the reference for freeze-dried samples by a constant offset δ (true freeze-dried concentration = reference + δ) and solved for the range of δ over which freeze-dried RMSE remained below oven-dried RMSE for the same milling treatment. We compared this range with the measured offset (n = 11 pairs, 95% CI from a paired t-test) in absolute terms and scaled by the relative offset. For calcium, which was not measured by IRMS, we scaled the relative offsets in carbon and nitrogen to mean observed calcium. This assumes a constant offset within a treatment and that ball-milled oven-dried cells carry no offset.

All spectral processing and trait predictions were performed with R v 4.4.1 (R Core Team, 2024). Analysis code was developed with assistance from a generative AI coding assistant (Claude Opus 5, Anthropic; used September 2026, Claude Sonnet 5.5, Anthropic; used October 2026) for code development, review, and debugging. All code was reviewed and executed by the authors; AI-suggested changes were accepted only after inspection and after the analysis reproduced prior results. The full analysis code, including its version history, is archived at (Frye *et al*., 2026b), and the quality control procedures applied to the input data are documented in Frye, (2026).

## RESULTS

### Variation of spectra between treatments

Across the full spectral range, the mean difference of reflectance between milling treatments ranged from 1.72–4.64 percent reflectance (Table 1). Across comparisons of milling treatments, the mean difference increased towards longer wavelengths with the average difference varying two to three-fold between the visible (400–700 nm) and SWIR2 (2001–2500 nm) regions. The deciduous broadleaf species tended to have greater differences between milling treatments compared to the needle and scale-leaved species (Table 1; Fig. 2). The highest mean reflectance differences were found between oven-dried fine and coarse mesh samples, with fine mesh sample reflectance having an average increase of 4.64 ± 0.98% reflectance. When compared to the coarse mesh, ball-milled samples had on average 2.92± 0.56% higher reflectance and the ball-milled treatment had 1.72± 0.91% lower reflectance compared to the fine mesh treatment. Because of practical challenges in milling needle-leaved samples to the finest grinds (these samples could only be milled through a 40-mesh screen versus the 60-mesh for broadleaves), particle size differs between growth forms at the fine mesh level (mean 137 µm in broadleaves versus 224 µm in needle-leaved material). This causes the broadleaf/needle-leaf contrasts in the milling effect to be confounded with treatment intensity. To account for this, we built a linear regression across the 21 trees with particle size measures (see below), regressing the particle size difference and a growth form term onto the fine – coarse spectral differences between the reduction in particle size (20-mesh minus fine, µm). The fine versus coarse spectral difference increased with the reduction in particle size (2.5% reflectance per 100 µm), and the growth-form term was not distinguishable from zero after account for it, although the interval was wide. Thus, after accounting for the reduction in particle size, broad- and needle-leave trees did not differ detectably in their spectral response to milling (a needle-leaf – broadleaf difference of -1.3 percentage points, 95% CI -5.6 to 2.9, compared with a raw difference of 4.9% among these 21 trees; 4.2% across all 61 trees; Table 1).

**Figure 2:**
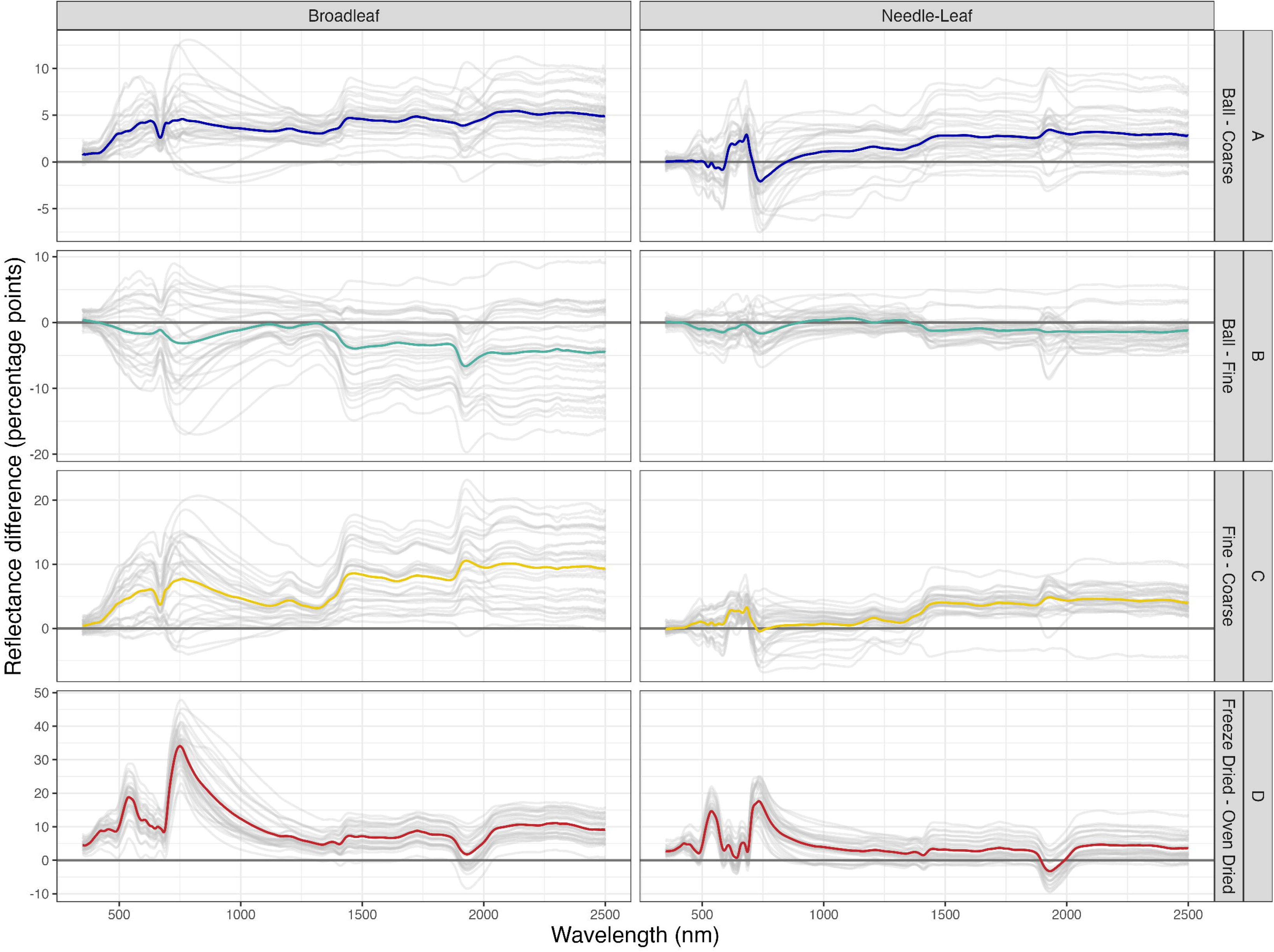
Finer milling and freeze-drying raised reflectance across the spectrum, most strongly in broadleaf species. Difference spectra (first treatment minus second; percentage points of reflectance; 350–2500 nm) for broadleaf (left, n = 30 trees) and needle- and scale-leaf species (right, n = 31 trees): (A) ball-milled minus 20-mesh; (B) ball-milled minus fine-mesh; and (C) fine-mesh minus 20-mesh. A–C use oven-dried material. (D) freeze-dried minus oven-dried, 20-mesh. Grey lines are individual trees (biological replicates), each the mean of all 36 pairwise differences between the six replicate scans (technical replicates) of the two treatments. Colored lines are the means across trees. Fine mesh is 60-mesh for broadleaf and 40-mesh for needle-leaf species. ***Alt Text Figure 2:*** *Grid of eight line graphs with four rows of treatment comparisons and two columns, broadleaf on the left and needle-leaf on the right. Each graph plots reflectance difference in percentage points against wavelength from 350 to 2500 nanometers, with grey lines for individual trees and one colored line for the mean. Row A, ball-milled minus coarse: the mean is positive, about 3 to 5 points in broadleaf and 0 to 3 in needle-leaf. Row B, ball-milled minus fine: the mean is near zero in needle-leaf and falls to about minus 5 in broadleaf beyond 1400 nanometers. Row C, fine minus coarse: the largest milling difference, rising to about 10 points beyond 1400 nanometers in broadleaf and about 4 in needle-leaf. Row D, freeze-dried minus oven-dried: sharp peaks near 550 and 750 nanometers, reaching about 34 points in broadleaf and 17 in needle-leaf, then declining through the infrared, with a dip near 1930 nanometers. In rows A to C, needle-leaf differences are smaller than broadleaf ones*.

**Table 1:** Mean reflectance difference between treatments by spectral region, in percentage points of reflectance. Each difference is the first treatment minus the second. Milling comparisons use oven-dried material; the drying comparison uses 20-mesh material. For each tree, the value is the mean of all 36 pairwise differences between the six replicate scans (technical replicates) of the two treatments. Values are means across trees (biological replicates) ± the half-width of the t-based 95% confidence interval. Coarse is 20-mesh; fine is 60-mesh for broadleaf and 40-mesh for needle- and scale-leaf species.

|  | **Ball - Coarse** |  |  | **Ball - Fine** |  |  | **Fine - Coarse** |  |  | **Freeze Dried - Oven Dried** |  |  |
|---|---|---|---|---|---|---|---|---|---|---|---|---|
| **Spectral Region** | **All Data<br>(n = 61)** | **Broadleaved<br>(n = 30)** | **Needle-Leaved<br>(n = 31)** | **All Data<br>(n = 61)** | **Broadleaved<br>(n = 30)** | **Needle-Leaved<br>(n = 31)** | **All Data<br>(n = 61)** | **Broadleaved<br>(n = 30)** | **Needle-Leaved<br>(n = 31)** | **All Data<br>(n = 61)** | **Broadleaved<br>(n = 30)** | **Needle-Leaved<br>(n = 31)** |
| Entire Spectral Range | 2.92 ± 0.56 | 4.08 ± 0.66 | 1.79 ± 0.72 | -1.72 ± 0.91 | -2.70 ± 1.74 | -0.77 ± 0.56 | 4.64 ± 0.98 | 6.78 ± 1.58 | 2.57 ± 0.65 | 7.15 ± 1.15 | 10.29 ± 1.32 | 4.12 ± 1.10 |
| Visible (400 nm-700 nm) | 1.78 ± 0.47 | 3.06 ± 0.53 | 0.55 ± 0.47 | -0.95 ± 0.68 | -1.13 ± 1.32 | -0.77 ± 0.49 | 2.73 ± 0.74 | 4.19 ± 1.25 | 1.32 ± 0.47 | 8.47 ± 1.10 | 11.44 ± 1.35 | 5.59 ± 0.95 |
| Near Infrared (701 nm-1500 nm) | 2.34 ± 0.63 | 3.76 ± 0.72 | 0.96 ± 0.79 | -0.89 ± 0.82 | -1.60 ± 1.57 | -0.20 ± 0.54 | 3.22 ± 0.96 | 5.36 ± 1.47 | 1.15 ± 0.75 | 8.83 ± 1.41 | 12.80 ± 1.69 | 5.00 ± 1.17 |
| Short Wave Infrared 1 (1501 nm-2000 nm) | 3.61 ± 0.58 | 4.41 ± 0.71 | 2.83 ± 0.86 | -2.51 ± 1.01 | -3.93 ± 1.85 | -1.15 ± 0.67 | 6.12 ± 1.05 | 8.34 ± 1.70 | 3.98 ± 0.74 | 4.16 ± 1.03 | 6.42 ± 1.32 | 1.97 ± 1.18 |
| Short Wave Infrared 2 (2001 nm-2500 nm) | 4.09 ± 0.68 | 5.19 ± 0.88 | 3.02 ± 0.92 | -2.91 ± 1.28 | -4.47 ± 2.41 | -1.40 ± 0.81 | 7.00 ± 1.29 | 9.65 ± 2.11 | 4.42 ± 0.87 | 6.97 ± 1.21 | 9.97 ± 1.40 | 4.07 ± 1.34 |

Differences in reflectance across milling treatments were not uniform and tended to increase around the following localized peaks: 626, 767, 1202, 1466, 1728, 1928, and 2428 nm (Figs. 2A–C). Note that the 2428 nm peaks could be an artifact of low signal-to-noise towards the ends of the spectral range. These increases are strongest in the comparison between coarse and finely milled samples but also appear in the same areas in the other milling comparisons.

Freeze-dried samples had higher reflectance than oven-dried samples (Fig. 2D). The average difference between all samples was 7.2%, with greater differences in broad-leaves compared to needle- and scale-leaved species (Table 1). Unlike the comparison of milling treatments, the greatest amounts of variation were in the visible and near infrared regions, reflecting the fact that oven-drying disproportionately degrades pigments. However, drying effects were less salient in the SWIR1 region (1501–2000 nm) compared to milling effects with a ratio of drying to milling variability of 0.68. The effects were equally impactful for SWIR2 (2001–2500 nm) with a ratio of 1.00. The maximum difference values found between drying treatments were 33.99% at 750 nm in broadleaf species and 17.62% at 733 nm in needle- and scale-leaf species. Localized peaks in the difference between oven-dried and freeze-dried spectra for broadleaves were near the following wavelengths: 456, 539, 750, 1187, 1449, 1727, 2297 nm (Fig. 2). These peaks also appeared in similar areas in the needle-leaf species, though the differences were more subdued.

Vector-normalization is a common pre-processing transformation in the remote sensing literature since it has been shown to reduce brightness differences caused by canopy architecture (Feilhauer *et al*., 2010). Because of this pre-processing step’s use in airborne models, it has also been applied to models with foliar powders (Wang *et al*., 2020). Separately, the near infrared spectroscopy field has long applied pre-processing transformations of standard normal variate (SNV) and multiple scatter correction with the explicit goal of reducing the effects of particle size differences (Geladi *et al*., 1985; Barnes *et al*., 1989; Dhanoa *et al*., 1994). We compared the differences between experimental treatments in the vector-normalized spectra (Supplementary Fig. S3) and found that spectral differences occurred in the same spectral regions as the raw measurements with minor differences in the near infrared region. We also compared the impacts that SNV and vector normalization have on spectra differences, finding that SNV was more effective in reducing particle size differences between samples compared to vector-normalization (Supplementary Table S1 and Supplementary Fig. S4). The milling share of variance fell from 24% (no transformation) to 17% with vector normalization and 5% with SNV. This suggests that researchers in the remote sensing field should consider using the SNV or similar corrections for foliar powders and reserve the use of vector-normalization for image-based models.

### Validation of spectrally predicted trait across treatments

When partial least squares regression coefficients (originally trained on oven-dried material ground through a rotary mill with a 20-mesh sieve) were applied to the vn-spectra across the milling and drying treatments, the oven-dried samples over-predicted nitrogen concentrations across all three milling treatments (Fig. 3A). The freeze-dried treatment over-predicted nitrogen at low concentrations and under-predicted at high concentrations. The drying treatments yielded NRMSE differences up to ∼7 percentage points, while the milling treatments had NRMSE differences of 2.6% for freeze-dried samples and 4.7% for oven-dried samples (Fig. 3A). The fine rotary treatment had the lowest difference in NRMSE performance between oven-dried and freeze-dried samples. Trends for calcium prediction were similar to nitrogen in that sample drying impacted prediction to a much greater extent compared to milling. Oven-dried samples tended to under-predict high calcium values to a greater degree than freeze-dried samples, and over-predicted low values to a lesser degree (Fig. 3B). Bland–Altman plots (Supplementary Fig. S5) showed that prediction error changed systematically with concentration, becoming more negative at high calcium in both drying treatments and, for nitrogen, increasing with concentration in oven-dried samples but decreasing in freeze-dried samples, consistent with the proportionality component.

**Figure 3:**
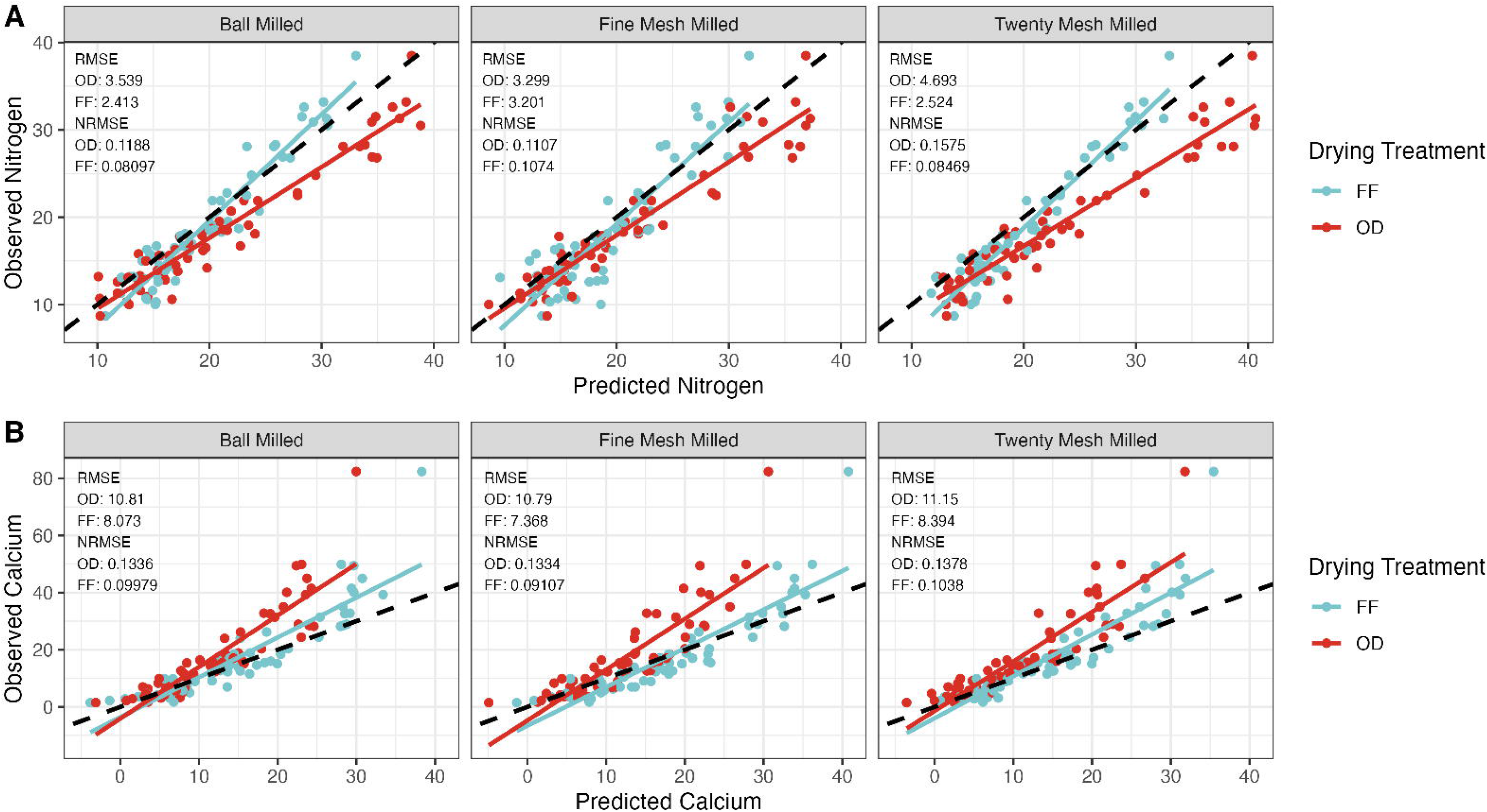
Freeze-dried powders gave lower prediction error than oven-dried powders for both traits in every milling treatment. Observed versus predicted (A) nitrogen and (B) calcium (mg g⁻¹), using the NEON PLSR coefficients of Wang et al. (2020). Each point is one tree (n = 60 per drying × milling combination). Predictions are the mean of six replicate scans (technical replicates). The observed value is a single assay of oven-dried, 20-mesh material, used as a common reference for all treatments. Solid lines are least-squares fits for each drying treatment; the dashed line is 1:1. RMSE is in mg g⁻¹; NRMSE is RMSE divided by the observed range. FF, freeze-dried; OD, oven-dried. ***Alt Text Figure 3:*** *Six scatterplots in two rows, nitrogen on top and calcium below, each with three columns for ball-milled, fine-mesh and twenty-mesh material. Observed concentration in milligrams per gram is plotted against predicted concentration, with points colored by drying treatment (light blue freeze-dried, red oven-dried), a fitted line for each drying treatment, and a dashed one-to-one line. For nitrogen, freeze-dried points lie close to the one-to-one line, while oven-dried points sit to its right, showing over-prediction that grows at higher concentrations. For calcium, predictions span about 0 to 40 while observed values reach 82 for one tree; the oven-dried fitted line is steeper than the one-to-one line, showing under-prediction of high values, and freeze-dried points lie closer to it. Each panel lists error values, which are lower for freeze-dried material in every panel*.

Coarsely milled, oven-dried samples yielded the highest NRMSE results for both traits. For nitrogen, freeze-dried ball-milled samples exhibited the lowest NRMSE value of 8.10% whereas the highest NRMSE value was 15.75% for coarsely milled, oven-dried samples. For calcium, freeze-dried finely milled samples exhibited the lowest NRMSE value of 9.11% and coarsely milled, oven-dried samples exhibited the highest NRMSE at 13.78%. In four of six treatments, nitrogen predictions had lower NRMSE values than estimates of calcium. Decomposing the error in the nitrogen predictions showed that freeze-dried advantage was largely a reduction in bias against the reference (0.58–1.25 mg g^-1^ bias for freeze-dried versus 2.18–3.82 mg g^-1^ for oven-dried) rather than a gain in precision (precision factor of the concordance correlation coefficient ρ=0.91–0.97 versus 0.95–0.97), although freeze-dried predictions were more compressed relative to the reference (regression slope of predicted on observed 0.71–0.76 versus 1.09–1.18; Supplementary Table S2). For calcium, the error decomposition showed that the freeze-dried advantage reflected a reduction in bias (−3.00 to +0.16 mg g⁻¹ for freeze-dried versus −6.79 to −5.21 mg g⁻¹ for oven-dried), and less proportional error (slope 0.55–0.60 versus 0.42–0.46) with similar precision (ρ = 0.90 versus 0.87–0.89; Supplementary Table S2).

### Variation of spectrally predicted traits in relation to inter- and intra-specific variation

The Bayesian ANOVA models demonstrate that for both calcium and nitrogen, different sample processing treatments impact spectral trait predictions (Fig. 4A). Freeze-drying was the most salient effect for predictions of calcium, with a posterior mean of 3.71 mg g^-1^ surrounded by a 95% credible interval (CI) of 2.92 – 4.48. This suggests that compared to oven-drying samples, freeze-drying is associated with higher calcium values. The ball mill and fine rotary mesh treatments also resulted in higher calcium predictions compared to the coarse mesh rotary mill. The ball mill posterior mean was 1.03 mg g^-1^ (CI: 0.25 – 1.80) and the fine mesh treatment posterior mean was 1.53 mg g^-1^ (CI: 0.74 – 2.32). While the credible interval of the interaction term for freeze-dried and ball mill samples overlapped 0, the posterior mean of the interaction term for freeze-drying and the fine rotary mill treatment was 1.60 mg g^-1^ (CI: 0.50 – 2.69). The species random-effect standard deviation (9.08 mg g⁻¹; 95% CI 5.92–14.23) was 3.8 times (CI 2.5–6.2) the SD of the six treatment-cell means. On average, 81% (65–92%) of the modelled variation in calcium was among species, 7% (3–15%) among trees within species, 5.4% (2.3– 10%) in the tree by treatment combination, 5.7% (2.4–10%) due to treatment, and under 1% scan-to-scan (Fig. 4B).

**Figure 4:**
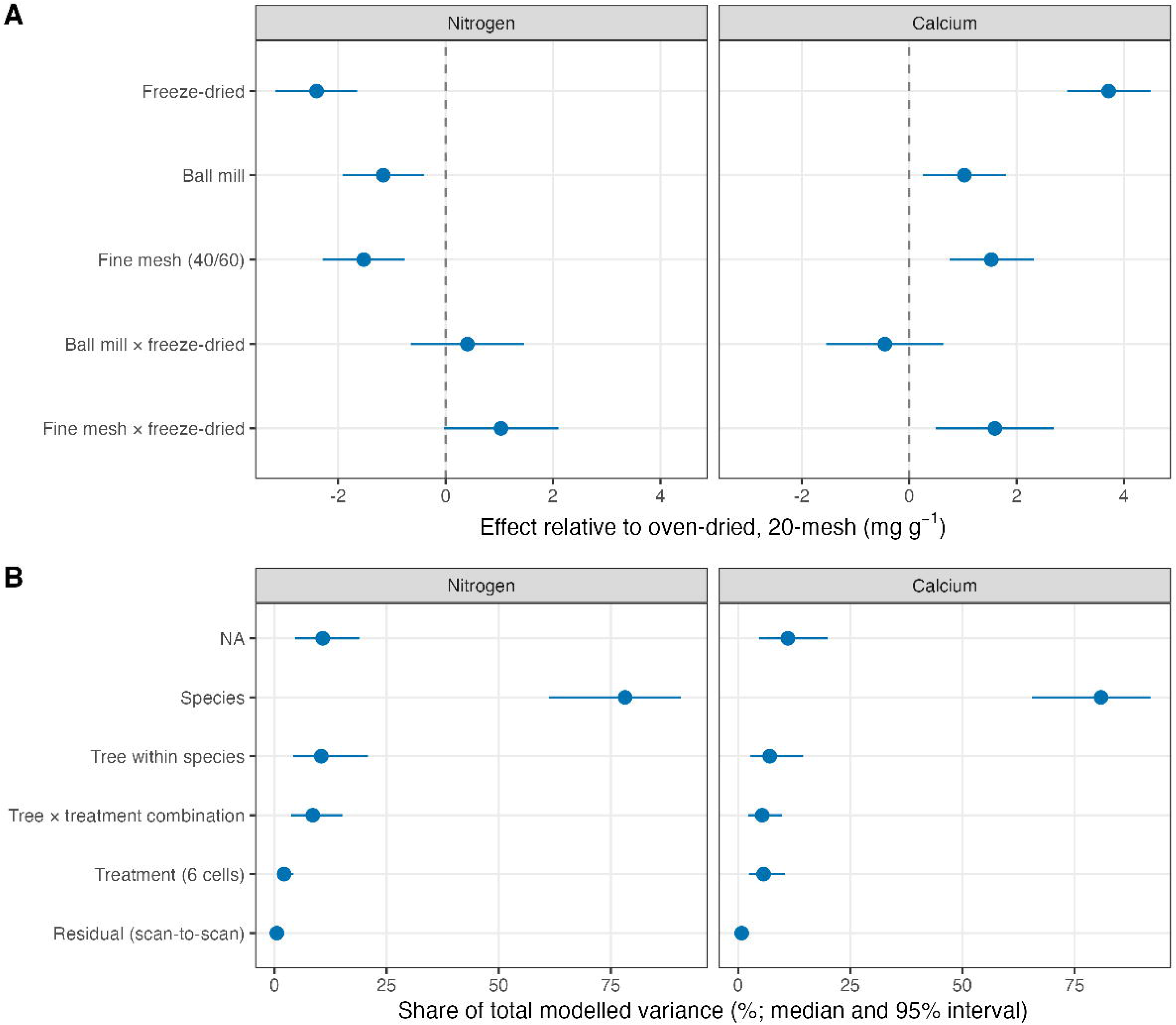
Drying had the largest processing effect on predicted traits, but species explained most of the variance. (A) Fixed-effect estimates (posterior median and 95% credible interval) from Bayesian multilevel models of spectrally predicted nitrogen and calcium. Effects are relative to oven-dried, 20-mesh material, in mg g⁻¹. (B) Share of total modelled variance (posterior median and 95% interval) among species, tree within species, the tree × treatment combination, treatment (the six drying × milling cell means) and residual scan-to-scan variation. Data are 61 trees of 11 species (biological replicates) × 6 treatments × 6 replicate scans (technical replicates), 2,196 scans in all. ***Alt Text Figure 4:*** *Two-part dot and interval chart with nitrogen on the left and calcium on the right. Part A shows estimated treatment effects in milligrams per gram, with a dashed line at zero. For nitrogen, freeze-drying, ball milling and fine-mesh milling are all negative, at about minus 2.4, minus 1.2 and minus 1.5, with intervals that exclude zero. Both interaction terms are positive, and their intervals touch or cross zero. For calcium, freeze-drying is about plus 3.7, ball milling about plus 1.0 and fine mesh about plus 1.5. The ball mill by freeze-drying interaction sits near zero and the fine mesh by freeze-drying interaction is about plus 1.6. Part B shows the share of variance for each level. Species dominates, at about 78 percent for nitrogen and 81 percent for calcium. Tree within species, tree by treatment, and treatment each account for about 11 percent or less, and scan-to-scan variation is near zero*.

Overall, the calcium model indicates that compared to calcium predictions using oven-dried samples milled through a 20-mesh screen, freeze-dried samples milled through a finer screen would result in significantly higher calcium predictions. The total of the freeze-drying term, the fine mesh term, and their interaction is 6.84 mg g^-1^. The average observed calcium from a sample that was oven-dried and milled through a 20-mesh rotary screen is 11.45 mg g^-1^. This suggests a 62.1% increase (CI 54.0 – 69.1) in calcium predictions given the highest differences in sample drying and milling, as calculated based on the sum of the freeze-dried, finely-milled, and interaction term divided by the observed mean.

Akin to the calcium models, freeze-drying was also the most salient treatment affecting nitrogen predictions with a mean estimate of -2.40 mg g^-1^ and 95% CI of -3.16 to -1.65. Both the ball milling and fine rotary mill treatments yielded lower nitrogen estimates with respective posterior mean estimates of -1.16 (CI: -1.92 to -0.39) and -1.52 (CI: -2.29 to -0.76). The interaction term for both ball-milling and freeze-drying overlapped 0 and the fine rotary milling and freeze-drying nearly overlapped 0 (lower bound −0.03). The trends in the random effect estimates were similar to the calcium model, with the magnitude of the standard deviation of the species random effect being higher than the treatment effects (6.92 with CI: 4.51 – 10.82). The individual and measurement replicate random effects were also nearly equivalent (2.46, CI: 1.96 – 3.08 and 2.20, CI: 2.03 – 2.39). Given that the average oven-dried and 20-mesh milled samples was 22.38 mg g^-1^, resulting in a 12.9% decrease (CI: -16.4 to -9.5) for a sample that was freeze-dried and finely-milled. For nitrogen, species accounted for 78% (61–91%) of the variation, trees within species 10% (4–21%), the tree × treatment combination 8.5% (3.7–15%), and treatment 2.1% (0.9–4.2%). The SD of the treatment-cell means was about half as large as for calcium (1.1 vs 2.3 mg g⁻¹), so processing mattered less for nitrogen predictions.

### Measured Particle Size Across Milling Treatments

The particle size distribution of a subset of oven-dried material was measured with a laser diffraction particle size analyzer to support interpretations of spectral measurements across different mill treatments. The milling treatments that exhibited the largest difference in particle size were the 20-mesh rotary milling and fine rotary milling treatments (Fig. 5). The transition from 20-mesh milled samples to finely milled samples provided a clear shift to smaller particle sizes as well as a smaller distribution range. Compared to 20-mesh milled samples, ball-milling shifted the distribution to smaller sizes, but had far less significant impact on overall particle size compared to finely milled samples. The average sample mean is 444.81 μm in 20 mesh samples, 333.53 μm in ball milled samples, and 178.21 μm in finely milled samples across all samples measured. The pooled standard deviations of the sample particle sizes measurements are 270.87 μm in 20-mesh milled samples, 242.11 μm in ball milled samples, and 119.73 μm in finely milled samples.

**Figure 5:**
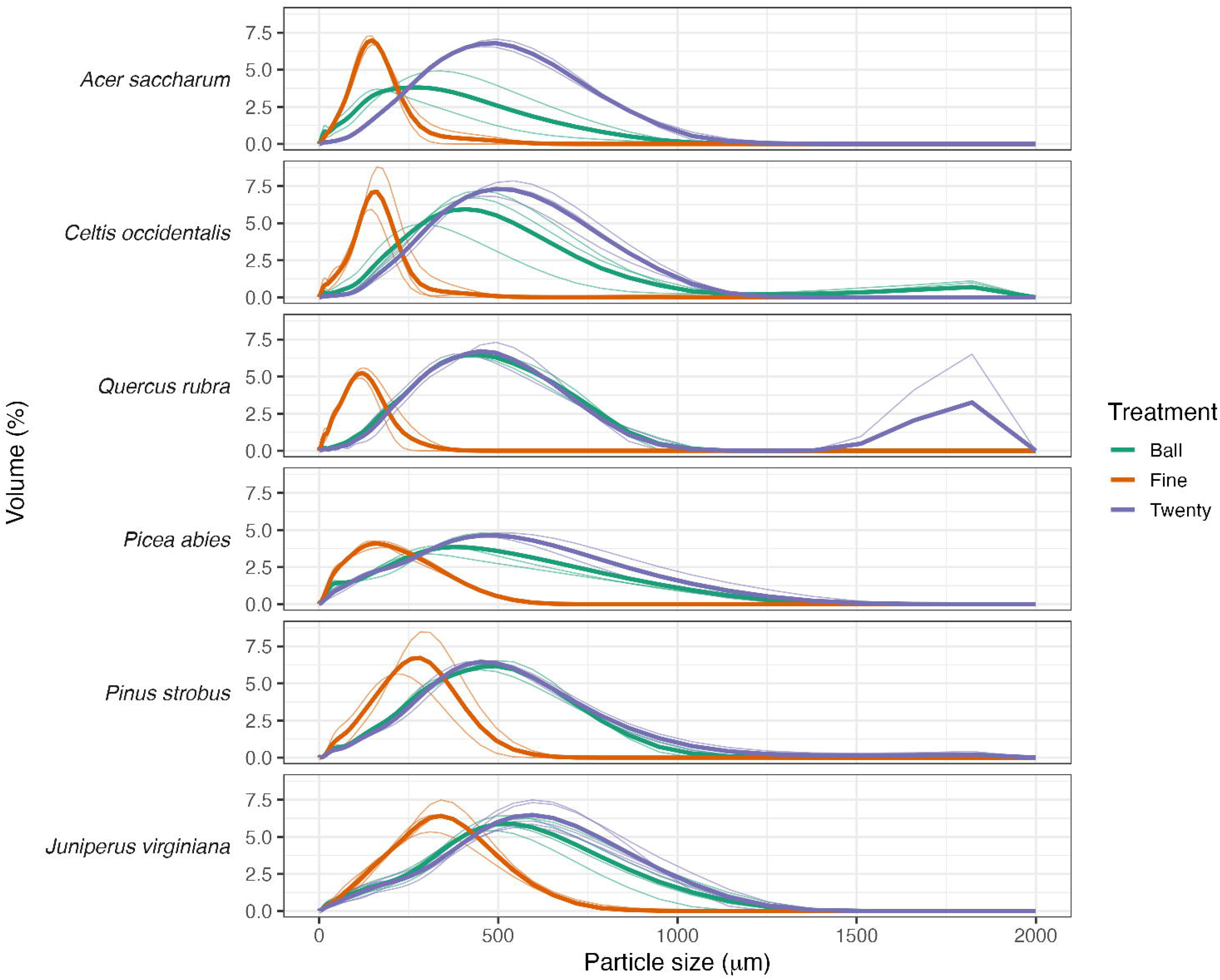
Fine-mesh milling produced the smallest and narrowest particle-size distributions, while ball milling overlapped 20-mesh in most species. Volume-weighted particle-size distributions of oven-dried material, measured by laser diffraction (LS 13 320), shown for each milling treatment and species. Thick lines show the species mean and thin lines show individual trees (n = 2–5 trees per species). Fine mesh is 60-mesh for broadleaf and 40-mesh for needle-leaf species. Five of the 22 measurements are not shown for clarity, but follow similar as those trends shown. ***Alt Text Figure 5:*** *Ridgeline plot of particle size distributions, one row per species: Acer saccharum, Celtis occidentalis, Quercus rubra, Picea abies, Pinus strobus and Juniperus virginiana. Each row shows three overlapping curves against particle size from 0 to 2000 micrometers: ball-milled in green, fine-mesh in orange and twenty-mesh in purple. In every species the fine-mesh curve is the narrowest and peaks at the smallest sizes, about 130 to 350 micrometers, with the needle-leaf species peaking at larger sizes than the broadleaf species. Twenty-mesh curves are broad, peak near 400 to 600 micrometers and tail beyond 1000 micrometers. Ball-milled curves closely overlap twenty-mesh in Celtis, Quercus and Pinus, sit between the other two in Picea and Juniperus, and peak at smaller sizes in Acer. Celtis ball-milled material has a small second peak near 1800 micrometers*.

## DISCUSSION

Our study indicates that sample processing affects spectral reflectance measurements and subsequent trait predictions. Flash-frozen and freeze-dried samples tended to have higher reflectance values compared to oven-dried material and yielded lower prediction errors with a partial least squares regression model that was initially trained using measurements of oven-dried material. While the effects of sample processing may be modest for robustly predicted foliar traits such as nitrogen, these processing differences can cause significant differences in predictions for traits such as calcium, which are less strongly predicted from reflectance spectra. Differences in milling procedure impacted spectral measurements as well, with the differences scaling with particle size. These results suggest that caution should be advised when merging data from studies with different sample processing procedures and that sample processing procedures should be carefully documented in the metadata of spectral datasets.

### Spectral reflectance differs between milling and drying treatments

More finely ground foliar material resulted in higher reflectance than coarsely ground material for both needle-leaf and broadleaf species (Fig. 2A–C). This result occurs because powders with smaller particles create more air to surface interfaces than larger ones, which in turn causes greater scattering. In other words, per a given distance within a powder, light will have more opportunity to reflect in a powder with smaller particles because those particles have greater specific area (Bull, 1991; Pasikatan *et al*., 2001). In addition to our study, this result has been shown in the near infrared spectroscopy literature (Devaux *et al*., 1995) as well as the field of soil science where reflectance is typically higher for finer soil particles (Sadeghi *et al*., 2018). Our direct measurements of particle size distribution for a subset of our samples support that the smallest particle size is produced by the fine mesh milling treatment compared to the 20-mesh rotary milling or ball milling. The contrasts between growth forms in milling effect are partly confounded with treatment intensity (60-vs. 40-mesh); particle-size reduction accounted for much of the gap, though with low power. We also note that the fine mesh milling treatment had a smaller standard deviation of particle size suggesting that this treatment produced a foliar powder that was more homogenized. However, this improved homogenization did not yield consistently strong gains in predicting the traits we assessed relative to the other milling approaches (Fig. 3). Frictional heating during ball-milling was not measured. Any heating that altered volatile or labile constituents could contribute to the milling effects we report, although ball-mill durations were short (2 min).

The effect of particle size tends to result in localized peaks of difference instead of a flat difference (Fig. 2). One mechanism driving these peaks is related to why reflectance is higher in finer powders— smaller particles not only create more opportunities for reflectance scatter but also absorption. Given the same substance but milled at two different particles sizes, the finer one will have stronger absorption features because of the increased interaction of light between particles (Bull, 1991). Given the position of the peaks, e.g., at 1940 nm, another driver of the reflectance difference between milling treatments may also be moisture uptake from ambient humidity (hygroscopic moisture). Devaux *et al*., (1995) measured a slight decrease in water content of more finely milled wheat flour and ground rape seed and Ben-Dor (1997) notes that there may be co-correlation between specific areas of particles and hygroscopic moisture. While it has been shown that additional drying of finer ground foliar materials increases trait prediction accuracy by reducing hygroscopic moisture (Ikoyi and Younge, 2020), Bull (1991) suggests that sample moisture is not the primary mechanism of these peaks and assigns the increased scattering from reduced particle size as the primary mechanism of localized moisture peaks.

Relative to milling comparisons, differences in reflectance between freeze-dried and oven-dried samples are an average of 1.54 times higher than the differences between finely milled and coarsely milled samples across the spectral range. This difference is largely linked to variability in the visible and near infrared regions. Milling treatments caused more variability in SWIR1 and equal variability SWIR2. Overall, freeze-dried samples exhibited higher reflectance across the spectrum compared to oven-dried samples (Fig. 2D). Within the visible region, this is expected because flash-freezing and then freeze-drying samples preserves pigments while the heat from oven-drying degrades them. One of the main peak differences of the drying treatment difference curve (Fig. 2D) is the large peak found at 750 nm. This peak is indicative of higher remnant concentrations of chlorophyll in freeze-dried samples compared to oven-dried samples (An-Erl *et al*., 2001). Higher reflectance in the near and short-wave infrared regions in freeze-dried materials is less clear but likely results from a similar process in which degradation of more reflective structural compounds is lower in freeze-dried samples. Proteins and starches have absorption features near localized peaks in spectral differences, e.g., 2068 nm and 2297 nm (Curran, 1989; Elvidge, 1990), suggesting that akin to pigments, they may be better preserved via freeze drying. While this does not likely change the elemental composition of the sample, it would change the overall reflectance. Lastly, freeze-dried samples likely had higher moisture contents than their oven-dried counterparts given localized absorptions peaks in the difference spectra (Fig. 2D). Freeze-dried samples were stored cold and scanned without a recorded equilibration step, so temperature or surface condensation could contribute to their stronger 1940 nm feature in addition to retained moisture within the powder; our data cannot separate these. The same applies to the isotope aliquots, which were weighed at ambient humidity, so moisture taken up during handling could also contribute to the lower carbon and nitrogen concentrations of the freeze-dried material (Supplementary Fig. S6).

### Impacts of sample processing on foliar trait predictions

Using calcium and nitrogen as example foliar traits, we found that sample handling has a meaningful impact on model results with freeze drying consistently yielding lower prediction errors. To a lesser degree, more finely milled samples also led to improvements in predictions except for freeze-dried nitrogen. These results were surprising because the models used to generate the spectral predictions were trained with data that was oven-dried and ground through a 20-mesh sieve (Wang *et al*., 2020). We originally anticipated that a model’s performance would favor new data that was processed more closely to the original methods of the model. However, we did not find this. Our decomposition of nitrogen model error suggests that prediction gains from freeze-drying largely reflected greater bias in the oven-dried predictions against the reference, not a gain in precision. For calcium, freeze-dried samples had lower bias and less proportional error (slope closer to 1), with similar precision. This could be because of higher spectral similarity with training data spectra, which we could not test. Note that the observed calcium range was set by one *Celtis occidentalis* (hackberry) tree (82 mg g⁻¹; the next highest was 50 mg g⁻¹); excluding it raised NRMSE (freeze-dried 10.5–12.1%, oven-dried 17.6–18.8%) and brought the regression slopes closer to one but did not change the ranking of treatments.

Because validation of spectral predictions was conducted against a single measured reference of oven dried and 20-mesh milled material, we conducted *post hoc* assessments of how sample handling could impact measured chemistry results and thus our subsequent validation. We found differences in the measured amounts carbon and nitrogen between samples that were freeze-dried versus oven-dried. Contrary to expectations dictated from carbon volatilization (Jones and O’Hara, 2016), the oven-dried material did not have lower carbon (which would cause higher nitrogen) when measured with IRMS (Supplementary Fig. S6). In fact, the freeze-dried material had lower carbon levels. Paired IRMS measurements (n = 11, ball-milled samples) showed freeze-dried material was lower than oven-dried material in both carbon (-15.9 mg g^-1^ or 3.2%) and nitrogen (-2.65 mg g^-1^ or 10%; 95% confidence interval -4.7 to -0.6; Supplementary Table S3). Given that the reference chemistry was oven-dried, we asked how large an offset in that reference would erase the freeze-dried NRMSE advantage using ball-milled pairs of samples (Supplementary Fig. S7). For nitrogen, the advantage persisted only for offsets between −2.1 and +3.2 mg g⁻¹ (ball milled) and between −0.3 and +2.0 mg g⁻¹ (fine mesh); the measured IRMS offset fell outside both ranges and inside the range for 20-mesh samples (−2.9 to +5.4), although its CI extended beyond the range in all three. We therefore interpret the nitrogen result as having closer agreement with an oven-dried reference and not necessarily a demonstrated gain in accuracy. For calcium the advantage persisted across offsets of −7.7 to +4.9 in all three milling treatments, and offsets scaled from carbon and nitrogen (−0.6 to −1.9 mg g⁻¹) lay well inside this range, so the calcium result was robust to plausible reference offsets. An explanation for the lower IRMS-measured carbon and nitrogen concentrations in freeze-dried samples is that those samples had higher moisture content, diluting the elemental concentrations. While carbon volatilization caused by vacuum pressure loss in the freeze-dried would also cause lower carbon, we would expect the nitrogen levels to increase, which we did not observe. Higher moisture content in our freeze-drying treatment is also supported by the difference around 1940 nm (a water absorption feature) in the spectral differences between oven-dried and freeze-dried samples (Fig. 2D). Despite the evidence that freeze-dried contained more moisture content, calcium predictions were higher based on the freeze-dried spectra; contrary to what would be expected if moisture was diluting elemental concentrations. This suggests that spectral predictions between treatments can be independent of the actual chemistry implying that the lower bias in the calcium models is not necessarily because of higher model accuracy. By contrast, the predicted nitrogen shift (freeze-dried – oven-dried; -3.1 mg g^-1^ for ball milled samples) was similar to the measured offset (-2.65 mg g^-1^), consistent with, but not demonstrating, a response to real compositional differences.

Lastly, we found that interspecific variation was greater than the variation attributable to sample processing (species standard deviation [SD] 6.0 times the treatment-cell SD for nitrogen and 3.8 for calcium; Fig. 4B). As expected from more global analyses, intra-specific variation was lower than inter-specific variation (Siefert *et al*., 2015). While sample processing had a lower impact on trait prediction compared to inter-specific variation for both traits, intraspecific variation was substantially closer to the variation caused by sample processing in the calcium and nitrogen models. Combining the treatment and specific treatment by tree effects yields that 10.7% (CI: 4.6–18.9) of the variation for nitrogen is attributable to treatment while the same calculation yields 11.1% (CI: 4.7 – 19.9) for calcium. These match or exceed the tree-within-species share (10.4% and 7.0%) with ratios of 1.03 and 1.57. The implication of these results is that researchers seeking to aggregate spectral datasets, especially across multiple species, may be able to do so as the amount of interspecific variation was much higher than that of the variation caused by differences in sample processing. However, caution should be advised for applications focused on more narrow windows of ecological variation, e.g., intra-specific variation or variation among closely related species. This is especially the case when modeling traits that have a weaker link to spectral features such as calcium.

### Practical applications

Our results demonstrate that sample handling meaningfully affects trait predictions from spectroscopy (Fig. 3). Freeze-drying improved agreement with the reference for Ca, an advantage that held across plausible reference offsets, but not demonstrably for N, and we found little evidence that finer milling improved predictions. Because freeze-drying requires considerably more labor than oven drying, and because accuracy cannot be separated from real treatment-induced chemical differences in our design, oven drying with consistent handling may be sufficient for many data collection efforts. However, depending on the research application, it may be important to halt leaf physiological processes and thus best preserve foliar chemistry, for which samples need to be flash-frozen in the field or relatively quickly in a laboratory once collected. Accessing and transporting liquid nitrogen in remote regions may be challenging, especially when large volumes of samples are being collected. While freezing of samples using dry ice is an alternative (used for pigment collection by the National Ecological Observatory Network (Weintraub, 2019)), we did not test this alternative, although we expect it to yield intermediate results because samples are not immediately frozen (compared to liquid N) and a limited amount of biological activity or degradation may occur. We also note that freeze dryers are more expensive than forced-air ovens and may be less effective at total moisture removal in powdered foliar materials (Adhamatika *et al*., 2021).

Whether larger spectral-trait datasets can be aggregated from sources with different sample handling procedures depends on the research objectives and still requires future work over different ecosystems and other types of foliar traits. Calibration transfer approaches similar to the ones used between spectroscopy instruments (Feudale *et al*., 2002; Yang *et al*., 2023) may need to be developed on datasets such as the one collected by this study.

## CONCLUSIONS

Spectral measurements of dried and homogenized foliar material are an important data source in ecological and agricultural studies that use these measurements to rapidly and cheaply assay a wide suite of biochemical plant traits. Spectroscopically predicted traits are often used as the basis for validating larger trait mapping efforts with air- and satellite-borne imaging spectroscopy (e.g., (Peterson *et al*., 1988; Bian *et al*., 2013; Serbin *et al*., 2014; Lepine *et al*., 2016; Wang *et al*., 2020; Liu *et al*., 2024) and there is an increasing interest to aggregate spectral-trait datasets into larger datasets akin existing plant trait repositories, e.g. TRY (Kattge *et al*., 2020). Despite the fact that spectroscopy of dried leaf powders has been used to assay plant properties for over fifty years since initial work of (Norris *et al*., 1976) little attention outside of the forage quality literature has been paid on how different processing methods affect measurements. Our study helps fill this gap by testing how different milling and drying methods affect spectral measurements and foliar trait predictions. While our study is limited to only two elemental foliar traits, we believe that our results are generalizable across plant traits with known spectral absorption features. We find that freeze drying and more finely milling samples generates higher reflectance values across the reflectance spectrum, yielding substantial impacts on the application of trait models. Researchers should pursue sample processing methods feasible for their studies while keeping in mind that these methodological decisions can impact results, and subsequently, how readily different spectral datasets can be integrated with one another.

## ACKNOWLEDGEMENTS

This study was supported by the following funding awarded to PAT: a National Science Foundation ASCEND Biology Integration Institute (BII) award DBI 2021898; a National Aeronautics and Space Administration (NASA) Jet Propulsion Laboratory subaward 1718573 and GSFC contract 80NSSC25PA477 in support of the AVUELO Campaign; and the NASA BioSCape award 80NSSC22K0905. This research was conducted at the UW-Madison Lakeshore Nature Preserve (permit 2024-LT#32) and supported by the University of Wisconsin-Madison Office of Sustainability. The authors gratefully acknowledge use of facilities and instrumentation in the UW-Madison Environmental Engineering Core Facility. The core facility is located within the Department of Civil & Environmental Engineering. This research was conducted using the Isotope Ratio Mass Spectrometry Core Facility at the Biotechnology Center of the University of Wisconsin–Madison, supported by the Shared Instrumentation Grant Program of the National Institutes of Health under award number 1S10OD036349-01. The content is solely the responsibility of the authors and does not necessarily represent the official views of the National Institutes of Health.

Declaration of generative-AI use. During the preparation of this manuscript the authors used Claude Opus 5 (Anthropic) to edit the abstract and cover letter for concision and clarity, and to check manuscript for internal consistency and factual accuracy against the cited sources and reported results. No text, data, figures, or citations were generated for inclusion without author verification; every flagged item was checked against the primary source by the authors. The authors reviewed and edited the output and take full responsibility for the content of the publication.

## AUTHOR CONTRIBUTIONS

Zachary Breuer - Data curation, visualization, validation, writing - original draft, writing - review and editing, formal analysis

Annabelle Majerus - Data curation, visualization, validation, writing - original draft, writing - review and editing, formal analysis

Henry A. Frye - Conceptualization, data curation, writing - original draft, writing - review and editing, formal analysis, supervision

Philip A. Townsend - Conceptualization, funding acquisition, supervision, writing - review and editing

## DATA AVAILABILITY

All data are available for download on Zenodo (10.5281/ZENODO.23504459) at Majerus *et al*., (2026). Code for data wrangling and analysis are available at (Frye *et al*., 2026b); (10.5281/zenodo.24143221). Code for data quality checks are available at Frye, (2026); (10.5281/ZENODO.24136458).

## ABBREVIATIONS

ICP-OES: inductively coupled plasma optical emission spectroscopy
IRMS: isotope-ratio mass spectrometry
MSE: mean squared error
NRMSE: normalized root mean squared error
RMSE: root mean squared error
SNV: standard normal variate
SWIR: short wave infrared
VN: vector-normalization/vector-normalized

## REFERENCES

Adhamatika A, Murtini ES, Sunarharum WB. 2021. The effect of leaf age and drying method on physico-chemical characteristics of pandan (Pandanus amaryllifolius Roxb.) leaves powder. IOP Conference Series: Earth and Environmental Science 733, 012073.

Alomar D, Fuchslocher R, Stockebrand S. 1999a. Effects of oven-or freeze-drying on chemical composition and NIR spectra of pasture silage. Animal Feed Science and Technology 80, 309–319.

Alomar D, Montero R, Fuchslocher R. 1999b. Effect of freezing and grinding method on near-infrared reflectance (NIR) spectra variation and chemical composition of fresh silage. Animal Feed Science and Technology 78, 57–63.

An-Erl KV, Liu C-F, Liu Y-J. 2001. Chlorophyll stability in spinach dehydrated by freeze-drying and controlled low-temperature vacuum dehydration. Food Research International 34, 167–175.

Asao S, Way DA, Turnbull MH, et al. 2025. Leaf nonstructural carbohydrate residence time, not concentration, correlates with leaf functional traits following the leaf economic spectrum in woody plants. New Phytologist 246, 1505–1519.

Barnes RJ, Dhanoa MS, Lister SJ. 1989. Standard Normal Variate Transformation and De-Trending of Near-Infrared Diffuse Reflectance Spectra. Applied Spectroscopy 43, 772–777.

Bellon-Maurel V, Fernandez-Ahumada E, Palagos B, Roger J-M, McBratney A. 2010. Critical review of chemometric indicators commonly used for assessing the quality of the prediction of soil attributes by NIR spectroscopy. TrAC Trends in Analytical Chemistry 29, 1073–1081.

Ben-Dor E. 1997. The reflectance spectra of organic matter in the visible near-infrared and short wave infrared region (400–2500 nm) during a controlled decomposition process. Remote Sensing of Environment 61, 1–15.

Bian M, Skidmore AK, Schlerf M, Wang T, Liu Y, Zeng R, Fei T. 2013. Predicting foliar biochemistry of tea (*Camellia sinensis*) using reflectance spectra measured at powder, leaf and canopy levels. ISPRS Journal of Photogrammetry and Remote Sensing 78, 148–156.

Bland MJ, Altman DG. 1986. Statstical Methods for Assessing Agreement Between Two Methods of Clinical Measurement. The Lancet 327, 307–310.

Blumstein M, Gersony J, Martínez-Vilalta J, Sala A. 2023. Global variation in nonstructural carbohydrate stores in response to climate. Global Change Biology 29, 1854–1869.

Bolster KL, Martin ME, Aber JD. 1996. Determination of carbon fraction and nitrogen concentration in tree foliage by near infrared reflectances: a comparison of statistical methods. Canadian Journal of Forest Research 26, 590–600.

Bull CR. 1991. Compensation for particle size effects in near infrared reflectance. The Analyst 116, 781.

Bürkner P-C. 2017. brms: An R Package for Bayesian Multilevel Models Using Stan. Journal of Statistical Software 80, 1–28.

Burnett AC, Anderson J, Davidson KJ, Ely KS, Lamour J, Li Q, Morrison BD, Yang D, Rogers A, Serbin SP. 2021. A best-practice guide to predicting plant traits from leaf-level hyperspectral data using partial least squares regression. Journal of Experimental Botany 72, 6175–6189.

Chadwick KD, Asner GP. 2016. Organismic-Scale Remote Sensing of Canopy Foliar Traits in Lowland Tropical Forests. Remote Sensing 8, 87.

Chlus A, Townsend PA. 2022. Characterizing seasonal variation in foliar biochemistry with airborne imaging spectroscopy. Remote Sensing of Environment 275, 113023.

Cloutis EA. 2015. Mineral and Rock Sample Database.

Couture JJ, Singh A, Rubert-Nason KF, Serbin SP, Lindroth RL, Townsend PA. 2016. Spectroscopic determination of ecologically relevant plant secondary metabolites. (M Davey, Ed.). Methods in Ecology and Evolution 7, 1402–1412.

Cryer J, Frye H, Zheng T, Townsend PA. 2026. Rotary milling.

Curran PJ. 1989. Remote sensing of foliar chemistry. Remote Sensing of Environment 30, 271–278.

Curran PJ, Dungan JL, Peterson DL. 2001. Estimating the foliar biochemical concentration of leaves with reflectance spectrometry. Remote Sensing of Environment 76, 349–359.

Devaux M-F, Nathier-Dufour N, Robert P, Bertrand D. 1995. Effects of Particle Size on the Near-Infrared Reflectance Spectra of Wheat and Rape Seed Meal Mixtures. Applied Spectroscopy 49, 84–91.

Dhanoa MS, Lister SJ, Sanderson R, Barnes RJ. 1994. The Link between Multiplicative Scatter Correction (MSC) and Standard Normal Variate (SNV) Transformations of NIR Spectra. Journal of Near Infrared Spectroscopy 2, 43–47.

Díaz S, Kattge J, Cornelissen JHC, et al. 2016. The global spectrum of plant form and function. Nature 529, 167–171.

Elvidge CD. 1990. Visible and near infrared reflectance characteristics of dry plant materials. International Journal of Remote Sensing 11, 1775–1795.

Eshel G, Levy GJ, Mingelgrin U, Singer MJ. 2004. Critical Evaluation of the Use of Laser Diffraction for Particle-Size Distribution Analysis. Soil Science Society of America Journal 68, 736–743.

Feilhauer H, Asner GP, Martin RE, Schmidtlein S. 2010. Brightness-normalized Partial Least Squares Regression for hyperspectral data. Journal of Quantitative Spectroscopy and Radiative Transfer 111, 1947–1957.

Feudale RN, Woody NA, Tan H, Myles AJ, Brown SD, Ferré J. 2002. Transfer of multivariate calibration models: a review. Chemometrics and Intelligent Laboratory Systems 64, 181–192.

Foley WJ, McIlwee A, Lawler I, Aragones L, Woolnough AP, Berding N. 1998. Ecological applications of near infrared reflectance spectroscopy - a tool for rapid, cost-effective prediction of the composition of plant and animal tissues and aspects of animal performance. Oecologia 116, 293–305.

Frye H. 2026. EnSpec/enspec-qc: Initial release.

Frye H, Breuer Z, Chlus A, et al. 2026a. Spectroscopy of dried and ground foliar material.

Frye H, Breuer Z, Majerus A, Townsend PA. 2026b. Analysis code for the effects of drying and milling treatments on estimates of foliar functional traits with VSWIR spectroscopy.

Frye H, Pribbenow M, Zheng T, Townsend PA. 2026c. Ball milling with paint shaker.

Geladi P, MacDougall D, Martens H. 1985. Linearization and Scatter-Correction for Near-Infrared Reflectance Spectra of Meat. Applied Spectroscopy 39, 491–500.

Green SJ, Brookson CB, Hardy NA, Crowder LB. 2022. Trait-based approaches to global change ecology: moving from description to prediction. Proceedings of the Royal Society B: Biological Sciences 289, 20220071.

Hawkesford MJ, Cakmak I, Coskun D, De Kok LJ, Lambers H, Schjoerring JK, White PJ. 2023. Functions of macronutrients. Marschner’s Mineral Nutrition of Plants. Elsevier, 201–281.

Ikoyi AY, Younge BA. 2020. Influence of forage particle size and residual moisture on near infrared reflectance spectroscopy (NIRS) calibration accuracy for macro-mineral determination. Animal Feed Science and Technology 270, 114674.

Jones DA, O’Hara KL. 2016. The influence of preparation method on measured carbon fractions in tree tissues. (M Ryan, Ed.). Tree Physiology 36, 1177–1189.

Kattge J, Bönisch G, Díaz S, et al. 2020. TRY plant trait database – enhanced coverage and open access. Global Change Biology 26, 119–188.

Kokaly RF, Asner GP, Ollinger SV, Martin ME, Wessman CA. 2009. Characterizing canopy biochemistry from imaging spectroscopy and its application to ecosystem studies. Remote Sensing of Environment 113, S78–S91.

Kothari S, Beauchamp-Rioux R, Blanchard F, et al. 2023. Predicting leaf traits across functional groups using reflectance spectroscopy. New Phytologist 238, 549–566.

Kothari S, Hobbie SE, Cavender-Bares J. 2024. Rapid estimates of leaf litter chemistry using reflectance spectroscopy. Canadian Journal of Forest Research 54, 978–991.

Kovach KR. 2023. SpectralPredictR. https://github.com/krkovach/SpectralPredictR?tab=readme-ov-file. Accessed September 2026.

Lepine LC, Ollinger SV, Ouimette AP, Martin ME. 2016. Examining spectral reflectance features related to foliar nitrogen in forests: Implications for broad-scale nitrogen mapping. Remote Sensing of Environment 173, 174–186.

Lin LI-K. 1989. A Concordance Correlation Coefficient to Evaluate Reproducibility. Biometrics 45, 255.

Liu N, Wagner Hokanson E, Hansen N, Townsend PA. 2023. Multi-year hyperspectral remote sensing of a comprehensive set of crop foliar nutrients in cranberries. ISPRS Journal of Photogrammetry and Remote Sensing 205, 135–146.

Liu S, Wang Z, Lin Z, Zhao Y, Yan Z, Zhang K, Visser M, Townsend PA, Wu J. 2024. Spectra-phenology integration for high-resolution, accurate, and scalable mapping of foliar functional traits using time-series Sentinel-2 data. Remote Sensing of Environment 305, 114082.

Majerus A, Breuer Z, Donnangelo G, Frye H, Kanski E, Pierce E, Vanden Heuvel C, Willems C, Townsend P. 2026. Supporting data for the effects of drying and milling treatments on estimates of foliar functional traits with VSWIR spectroscopy: leaf scans, processed spectra and replicate-pair differences.

Martin M, Plourde L, Ollinger S, Smith M-L, McNeil B. 2008. A generalizable method for remote sensing of canopy nitrogen across a wide range of forest ecosystems. Remote Sensing of Environment 112, 3511–3519.

Matson P, Johnson L, Billow C, Miller J, Pu R. 1994. Seasonal Patterns and Remote Spectral Estimation of Canopy Chemistry Across the Oregon Transect. Ecological Applications 4.

Mayland HF. 1968. Effect of Drying Methods on Losses of Carbon, Nitrogen and Dry Matter from Alfalfa. Agronomy Journal 60, 658–659.

Myers TL, Brauer CS, Su Y-F, Blake TA, Tonkyn RG, Ertel AB, Johnson TJ, Richardson RL. 2015. Quantitative reflectance spectra of solid powders as a function of particle size. Applied Optics 54, 4863–4875.

Norris KH, Barnes RF, Moore JE, Shenk JS. 1976. Predicting Forage Quality by Infrared Replectance Spectroscopy. Journal of Animal Science 43, 889–897.

Park RS, Agnew RE, Kilpatrick DJ. 2002. The effect of freezing and thawing on grass silage quality predictions based on near infrared reflectance spectroscopy. Animal Feed Science and Technology 102, 151–167.

Pasikatan MC, Steele JL, Spillman CK, Haque E. 2001. Near Infrared Reflectance Spectroscopy for Online Particle Size Analysis of Powders and Ground Materials. Journal of Near Infrared Spectroscopy 9, 153–164.

Peterson DL, Aber JD, Matson PA, Card DH, Swanberg N, Wessman C, Spanner M. 1988. Remote sensing of forest canopy and leaf biochemical contents. Remote Sensing of Environment 24, 85–108.

Popp M, Lied W, Meyer AJ, Richter A, Schiller P, Schwitte H. 1996. Sample preservation for determination of organic compounds: microwave versus freeze-drying. Journal of Experimental Botany 47, 1469–1473.

Prananto J, Minasny B, Weaver T. 2020. Near infrared (NIR) spectroscopy as a rapid and cost-effective method for nutrient analysis of plant leaf tissues. Advances in Agronomy.

Quentin AG, Pinkard EA, Ryan MG, et al. 2015. Non-structural carbohydrates in woody plants compared among laboratories. (M Mencuccini, Ed.). Tree Physiology, tpv073.

R Core Team. 2024. R: A Language and Environment for Statistical Computing. Vienna, Austria: R Foundation for Statistical Computing.

Robeson SM, Willmott CJ. 2023. Decomposition of the mean absolute error (MAE) into systematic and unsystematic components. (F Zama, Ed.). PLOS ONE 18, e0279774.

Sadeghi M, Babaeian E, Tuller M, Jones SB. 2018. Particle size effects on soil reflectance explained by an analytical radiative transfer model. Remote Sensing of Environment 210, 375–386.

Serbin SP, Singh A, McNeil BE, Kingdon CC, Townsend PA. 2014. Spectroscopic determination of leaf morphological and biochemical traits for northern temperate and boreal tree species. Ecological Applications 24, 1651–1669.

Siefert A, Violle C, Chalmandrier L, et al. 2015. A global meta-analysis of the relative extent of intraspecific trait variation in plant communities. (J Chase, Ed.). Ecology Letters 18, 1406–1419.

Singh A, Serbin SP, McNeil BE, Kingdon CC, Townsend PA. 2015. Imaging spectroscopy algorithms for mapping canopy foliar chemical and morphological traits and their uncertainties. Ecological Applications 25, 2180–2197.

Smith D. 1973. Influence of Drying and Storage Conditions on Nonstructural Carbohydrate Analysis of Herbage Tissue—a Review. Grass and Forage Science 28, 129–134.

Thompson K, Parkinson JA, Band SR, Spencer RE. 1997. A comparative study of leaf nutrient concentrations in a regional herbaceous flora. New Phytologist 136, 679–689.

Townsend PA, Foster JR, Chastain RA, Currie WS. 2003. Application of imaging spectroscopy to mapping canopy nitrogen in the forests of the central Appalachian Mountains using Hyperion and AVIRIS. IEEE Transactions on Geoscience and Remote Sensing 41, 1347–1354.

Wagner EP, Merz J, Townsend PA. 2019. EcoSIS: A Spectral Library and the Tools to Use It. AGU Fall Meeting Abstracts. AGU Fall Meeting Abstracts. B11F-2396.

Wang Z, Chlus A, Geygan R, Ye Z, Zheng T, Singh A, Couture JJ, Cavender-Bares J, Kruger EL, Townsend PA. 2020. Foliar functional traits from imaging spectroscopy across biomes in eastern North America. New Phytologist 228, 494–511.

Watanabe T, Broadley MR, Jansen S, White PJ, Takada J, Satake K, Takamatsu T, Tuah SJ, Osaki M. 2007. Evolutionary control of leaf element composition in plants. New Phytologist 174, 516–523.

Weintraub S. 2019. TOS Protocol and Procedure: Canopy Foliar Sampling.

Wessman CA, Aber JD, Peterson DL, Melillo JM. 1988. Remote sensing of canopy chemistry and nitrogen cycling in temperate forest ecosystems. Nature 335, 154–156.

Willmott CJ. 1981. On the Validation of Models. Physical Geography 2, 184–194.

Wold S, Sjöström M, Eriksson L. 2001. PLS-regression: a basic tool of chemometrics. Chemometrics and Intelligent Laboratory Systems 58, 109–130.

Wright IJ, Reich PB, Westoby M, et al. 2004. The worldwide leaf economics spectrum. Nature 428, 821–827.

Yang X, Cherney J, Casler M, Berzaghi P. 2023. Forage calibration transfer from laboratory to portable near infrared spectrometers. Journal of Near Infrared Spectroscopy 31, 126–140.

Youngflesh C. 2018. MCMCvis: Tools to Visualize, Manipulate, and Summarize MCMC Output. Journal of Open Source Software 3, 640.

